# FGFs are orchestra conductors of Shh-dependent oligodendroglial fate specification in the ventral spinal cord

**DOI:** 10.1101/233775

**Authors:** Marie-Amélie Farreny, Eric Agius, Sophie Bel-Vialar, Nathalie Escalas, Nagham Khouri-Farah, Chadi Soukkarieh, Fabienne Pituello, Philippe Cochard, Cathy Soula

## Abstract

Most oligodendrocytes of the spinal cord originate from ventral progenitor cells of the pMN domain, characterized by expression of the transcription factor Olig2. A minority of oligodendrocytes is also recognized to emerge from dorsal progenitors during fetal development. The prevailing view is that generation of ventral oligodendrocytes depends on Sonic hedgehog (Shh) while dorsal oligodendrocytes develop under the influence of Fibroblast Growth Factors (FGFs). Using the well-established model of the chicken embryo, we evidence that ventral spinal progenitor cells activate FGF signaling at the onset of oligodendrocyte precursor cell (OPC) generation, as do they dorsal counterpart. Inhibition of FGF receptors at that time appears sufficient to prevent generation of ventral OPCs, highlighting that, in addition to Shh, FGF signaling is required also for generation of ventral OPCs. We further reveal an unsuspected interplay between Shh and FGF signaling by showing that FGFs serve dual essential functions in ventral OPC specification. FGFs are responsible for timely induction of a secondary Shh signaling center, the lateral floor plate, a crucial step to create the burst of Shh required for OPC specification. At the same time, FGFs prevent down-regulation of Olig2 in pMN progenitor cells as these cells receive higher threshold of the Shh signal. Finally, we bring arguments favoring a key role of newly differentiated neurons acting as providers of the FGF signal required to trigger OPC generation in the ventral spinal cord.

## Introduction

Oligodendrocytes, the myelin-forming cells of the vertebrate central nervous system (CNS), differentiate from oligodendrocyte precursor cells (OPCs) generated during development. OPCs are specified at embryonic stages and are known to originate from multiple progenitor domains of the ventricular zone (Rowitch and Kriegstein, 2010). In the developing spinal cord, two spatially and temporally distinct sources of OPCs have been evidenced. An initial ventral source, producing most OPCs of the spinal cord, emanates from a subset of ventral neural progenitors populating the so called pMN domain (Rowitch, 2004). A later dorsal source that produces a minority of OPCs, starting at fetal stages, takes place in more dorsal progenitor domains (Cai et al., 2005; Fogarty et al., 2005; Vallstedt et al., 2005). The importance of extracellular signaling factors in defining both the dorso-ventral position of OPC sources and the precise timing of OPC specification are widely recognized. Induction of ventral OPCs depends on the morphogenetic activity of Sonic Hedgehog (Shh) produced by ventral medial cells of the developing spinal cord (Danesin and Soula, 2017). During this first wave of production, the restriction of OPC generation to ventral progenitors is also controlled by the repressive activity of the dorsalizing factors BMPs and Wnts which at that time prevent OPC generation from dorsally located neural progenitor cells (Mekki-Dauriac et al., 2002; Miller et al., 2004). In contrast, the second phase of OPC generation from dorsal progenitor cells occurs independently of Shh and has been proposed instead to depend on Fibroblast growth factors (FGFs) that have been proposed to act by counteracting BMP signaling (Chandran et al., 2003; Kessaris et al., 2004; Vallstedt et al., 2005; Bilican et al., 2008). Therefore, it is currently assumed that commitment of ventral and dorsal spinal cord OPCs depends on Shh and FGFs, respectively. Similarly, multiple spatial and temporal waves of OPC generation have been reported to occur in the developing brain. The first wave of OPCs occurs in the ventral forebrain and requires Shh activity (Alberta et al., 2001; Nery et al., 2001; Spassky et al., 2001; Tekki-Kessaris et al., 2001b, a). However, evidences has further emerged that generation of ventral OPCs in the mouse forebrain also depends on the FGF signaling pathway (Furusho et al., 2011). A similar observation has been made in the zebrafish ventral rhombencephalon where FGF signaling has been shown to cooperate with Shh to control expression of Olig2 (Esain et al., 2010). Therefore, contrasting with the proposed model for the spinal cord, ventral OPC generation in the developing brain depends on a Shh and FGF interplay, whose precise mechanism remains, however, to be elucidated.

Although a number of studies established a framework for understanding mechanisms controlling Shh-dependent OPC induction in the ventral spinal cord, a number of questions remain. A prerequisite for ventral OPC generation is establishment of the pMN domain characterized by expression of Olig2, an obligatory transcription factor for oligodendrocyte development. This domain initially forms in the ventral neural tube in response to Shh secreted from the notochord and medial floor plate (MFP) cells (Dessaud et al., 2008). Once established, the pMN domain produces motor neurons (MNs) and start generating OPCs only after completion of MN generation (Bergles and Richardson, 2015). Shh is still necessary at these late stages for OPC induction (Orentas et al., 1999; Soula et al., 2001; Agius et al., 2004; Park et al., 2004). Significantly, Olig2 progenitor cells must not only receive Shh but must be submitted to a high threshold Shh signal to change their fate and generate OPCs (Soula et al., 2001; Danesin et al., 2006). Accordingly, a temporal rise of Shh signaling occurs in the ventral spinal cord at the time of OPC specification (Danesin et al., 2006; Touahri et al., 2012; Al Oustah et al., 2014). This results in up-regulation of the homeodomain protein Nkx2.2 within Olig2-expressing progenitor cells that finally ends up with establishment of a new domain, named the p* domain, populated by Nkx2.2 and Olig2-coexpressing cells (Fig. 1A)(Qi et al., 2001; Zhou et al., 2001; Fu et al., 2002; Agius et al., 2004; Touahri et al., 2012). During patterning process of the neural tube, activation of Nkx2.2 in ventral-most progenitor cells is known to repress Olig2 expression, an essential function allowing formation of the two distinct p3 (Nkx2.2) and pMN (Olig2) domains (Jeong and McMahon, 2005; Stamataki et al., 2005; Dessaud et al., 2007). However, at the time of OPC specification, activation of Nkx2.2 in pMN cells no longer represses Olig2 and their coexpression drives progenitor cells to the OPC fate (Zhou et al., 2001; Sun et al., 2003). Thus, the Shh signal must not only remain active over development but must also be temporally strengthened to modify the transcriptional status of Olig2 progenitor cells. Changes in both Shh source cell identity and position, that undergo significant transformations over time, have been proposed to underlie timely activation of Shh signaling (Al Oustah et al., 2014; Danesin and Soula, 2017). In particular, a secondary source of Shh, named the lateral floor plate (LFP), forms shortly before OPC induction by up-regulation of Shh expression in progenitor cells of the p3 domain, thereby bringing a source of Shh in closer proximity to the pMN domain (Fig. 1A) (Charrier et al., 2002; Schäfer et al., 2007; Al Oustah et al., 2014; Jiang et al., 2017). However, how LFP induction is controlled over time and why Nkx2.2 activation no more represses Olig2 expression in cells of the pMN domain at initiation of OPC generation are still open questions.

**Figure 1:**
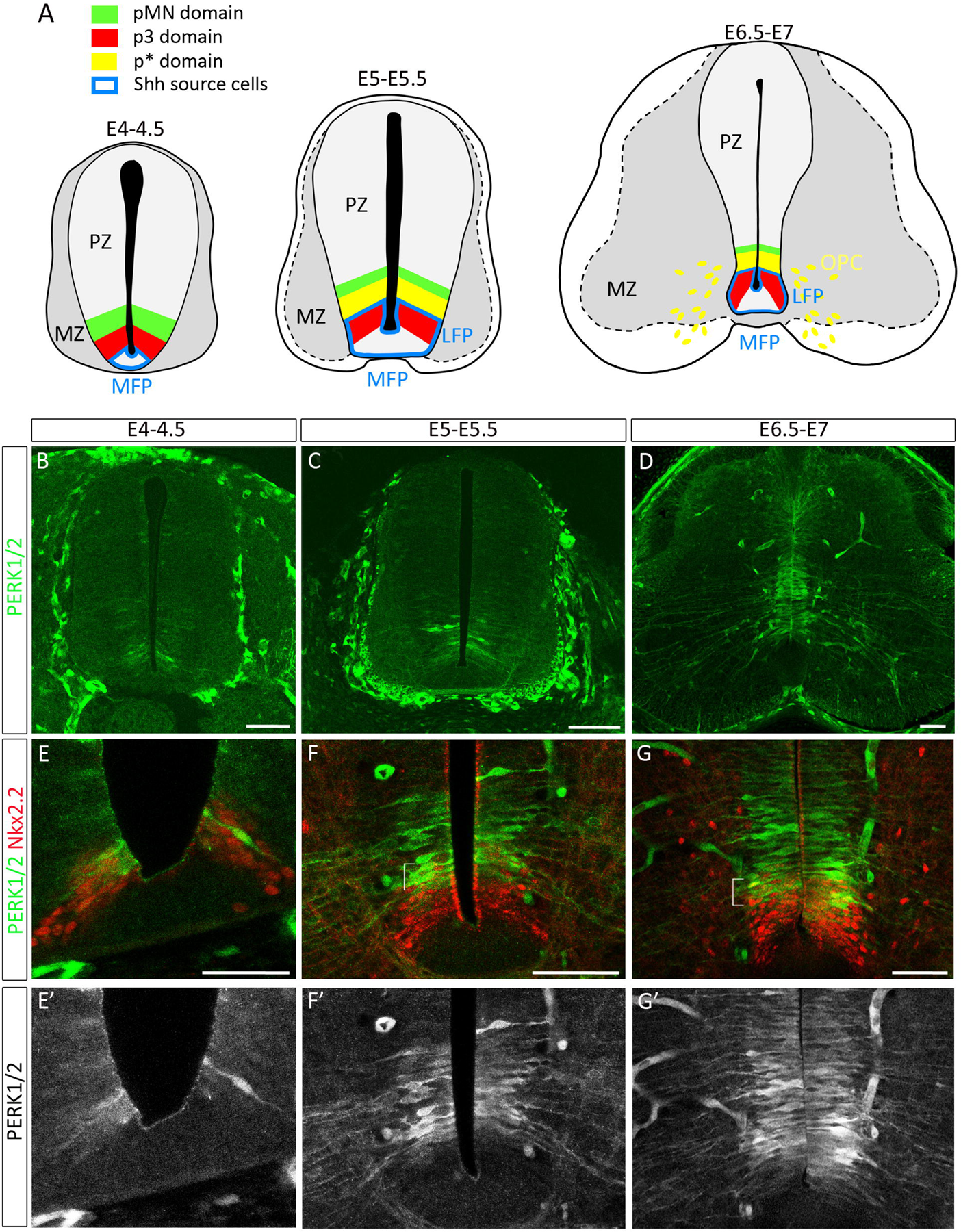
Activation of ERK1/2 in ventral spinal cord progenitor cells correlates with the time of OPC specification. **A**: Schematic of medial floor plate (MFP), lateral floor plate (LFP) and neural domain organization over development of the spinal cord, dorsal on top in all figures. At E4-E4.5, Shh expression (blue line) is restricted to MFP cells while Nkx2.2 and Olig2 are expressed in the adjacent non overlapping p3 (red) and pMN (green) domains, respectively. Following LFP formation (E5-E5.5), i.e. activation of Shh expression in the former p3 domain, progenitor cells of the pMN domain up-regulate Nkx2.2 leading to formation of the p* domain populated by Olig2 and Nkx2.2 coexpressing progenitor cells (yellow). From E6, Nkx2.2/Olig2 coexpressing OPCs (yellow) migrate from the p* domain to progressively populate the entire spinal cord. PZ = progenitor zone, MZ = marginal zone. **B-G**: Temporal profile of P-ERK1/2 on transverse sections of chicken spinal cord. Note the progressive ventral to dorsal activation of ERK1/2 in the ventral progenitor zone. **E-G**: Double detection of P-ERK1/2 (green) and Nkx2.2 (red) viewed at higher magnification of the ventral progenitor zone. Note high level of the P-ERK1/2 signal in the dorsal-most Nkx2.2-expressing cells (p* domain) at E5.5-E6 and 6.5-E7 (brackets in E and G). Scale bars = 50μm.

In this study, we bring answers to these issues by highlighting the key role of FGF signaling in controlling OPC induction also in the ventral spinal cord. Using chicken as a well-established model of OPC development, we provide evidence for a dual role of FGF signaling acting upstream of Shh to promote induction of the LFP but also together with Shh to ensure maintenance of Olig2 expression in pMN progenitor cells as they activate Nkx2.2 in response to high threshold Shh signal.

## Materials and Methods

Fertilized White Leghorn chicken eggs, obtained from a commercial source, were incubated at 38°C until they reached the appropriate stages (Hamburger and Hamilton, 1992). Animal related procedures were performed according to the EC guiding principles (86/609/CEE), French Decree no. 97/748 and the recommendations of the CNRS.

### Isolation and culture of spinal cord explants

Flat-mount preparations of spinal cord explants were cultivated using an organotypic culture system previously described (Agius et al., 2010). Briefly, E4 spinal cord explants, isolated from surrounding tissues, were opened along the dorsal midline and flattened on a nitrocellulose membrane (Sartorius) with the neural progenitor cells up and grown as organotypic cultures in DMEM (Invitrogen) supplemented with 10% FCS (Sigma). When indicated, the FGFR inhibitors SU5402 (Mohammadi et al., 1997) and PD166866 (Panek et al., 1998) or the Shh signaling inhibitor cyclopamine (Incardona et al., 1998) were added to the culture medium at the time of plating or after 24 hours in culture. The working concentrations of inhibitors were as follows: SU5402 20μM (Calbiochem); PD166866 2μM (Calbiochem); cyclopamin 5mM (Enzo life Sciences). Inhibitors were solubilized and diluted in DMSO that was added to control cultures. Flat-mounted explants were fixed at appropriate intervals in 3.7% formaldehyde/PBS, sectioned at 60-80μm using a vibratome (Microm) and processed for immunostaining or in situ hybridization.

### Tissue preparation

Chicken embryos or flat-mounted explants were fixed in 4% paraformaldehyde (PFA) in PBS. Embryos and explants were fixed overnight at 4°C or 1 hour at room temperature, respectively, for immunodetection analysis. When intended to in situ hybridization, an overnight fixation (4°C) was performed followed by dehydratation and storage in 100% ethanol (-20°C). Tissues were then sectioned at 60–80μm using a vibratome (Microm). In all experiments, sections were performed at the brachial level.

### Immunostaining

For immunofluorescence analysis, chicken tissues were processed as previously described (Agius et al., 2004; Danesin et al., 2006). Primary antibodies were applied overnight at 4°C. Secondary antibodies were applied at room temperature for 1 hour. The antibodies used were as follows: rabbit anti Phospho-p44/42 MAPK (PERK) 1/200 (Cell Signaling); rabbit anti-Olig2 1/500 (Chemicon); mouse monoclonal O4 antibody (O4 mAb), 1/4 (culture supernatant obtained from O4 hybridoma cells, a gift from R. Bansal); mouse anti-Nkx2.2 1/4 (DSHN) (Ericson et al., 1996); rabbit anti-clived Caspase 3 1/400 (Cell Signaling), mouse anti-BrdU 1/2000 (DHSB). Alexa 488, 555 or 647 goat anti-rabbit and goat anti-mouse (1/500) were from Molecular Probes. For BrdU staining, spinal cord explants were incubated for 2 hours with BrdU (0.15μg/μl, Sigma), 24, 32 or 48 hours after plating. Tissues were then fixed for 4 hours and processed for BrdU immunostaining.

### In situ hybridization

In situ hybridization (ISH) was performed on vibratome sections of chicken embryos and spinal cord explants either by hand or automatically (InsituPro, Intavis) using the whole-mount ISH protocol previously described (Braquart-Varnier et al., 2004; Danesin et al., 2006). Digoxigenin (DIG)-labeled sense and antisense RNA probes were synthesized using T3 and T7 polymerases. RNA labeled probes were detected by an alkaline-phosphatase coupled antibody (Roche Diagnostics) and NBT/BCIP (nitroblue tetrazolium/5-bromo-4-chloro-3-indolyl phosphate) was used as a chromogenic substrate for alkaline phosphatase (Boehringer, Mannheim). The following RNA probes were used: *spry1 and spry2, fgfr1, fgfr2* and *fgfr4* (provided by K. Storey); *mkp3* and *fgf8* (provided S Martinez), *Shh* (provided by C. Tabin). Counterstaining of Nkx2.2 was performed after color development following a post-fixation step in 4% PFA for 1 hour.

### Electroporation

Expression constructs were cloned into either the pCAG-IRES-GFP (Addgene) for the truncated FGF receptor (dnFGFR) (Amaya et al., 1991) or the pCMV vector for the chimeric protein FGF8b-GFP (Pombero et al., 2011). To allow cell body detection of electroporated cells, the pCMV FGF8b-GFP vector was co-electroporated with the empty pCIG vector (a gift from A. McMahon) used at 0.5μg/μl In ovo electroporation in E1.5 neural tube was performed as described previously (Itasaki et al., 1999). Briefly, the FGF8 and/or Shh constructs were injected at 1μg/μl in the rostral neural tube using a glass pipette. Electrodes (Nepa Gene Corporation) were positioned on each side of the neural tube and four pulses of 20 V (Intracel, TSS10) were applied to trigger unilateral entry of the DNA into the neural tube, the non-transfected half constituting an internal control. Electroporation of E4 spinal cord was performed ex ovo. The dnFGFR expression vector was used at 1μg/μl Controls were performed with pCAG-IRES-GFP vector alone. Embryos were harvested and isolated in a Petri dish with the dorsal side up, and DNA solution was injected into the lumen of the spinal cord as previously described (Danesin et al., 2006; Touahri et al., 2012). Electrodes were positioned on each side of the brachial region of the spinal cord, the positive electrode being placed more ventrally than the negative one, allowing satisfactory electroporation of ventral regions. Ten pulses of 25 V were applied and spinal cord was further dissected and grown in organotypic culture as above.

### Experimental Design and Statistical Analysis

Fluorescence photomicrographs were collected with Leica SP5 and Zeiss 710 confocal microscopes. Images of ISHs were collected with Nikon digital camera DXM1200C and a Nikon eclipse 80i microscope. Images were processed using Adobe Photoshop CS2. Unless otherwise stated in figure legends, provided data are the average of three embryos or explants (n) per condition from at least two independent experiments. Cell counting was performed on at least 5 tissue slices per chicken explant. For each tissue slice (60–80 μm), at least 4 optical sections were acquired at 6 μm intervals and cells were counted in each optical section. Quantifications are expressed as the mean number of cells (mean±s.e.m) in an optical section of hemi-explants. Statistical analyses were performed using the Mann–Whitney U test. Significance was determined at p < 0.05. p values are indicated in figure legends or in text when quantifications are not included in figures.

## Results

### MAPK signaling is activated at initiation of OPC commitment in the ventral spinal cord

Previous studies have reported that FGFs can induce production of OPCs from dorsal spinal cord and cerebral cortex progenitor cells (Chandran et al., 2003; Gabay et al., 2003; Kessaris et al., 2004; Cai et al., 2005; Abematsu et al., 2006; Naruse et al., 2006; Bilican et al., 2008). This inductive property has been attributed to robust activation of the MAPK signaling pathway (Chandran et al., 2003; Kessaris et al., 2004; Bilican et al., 2008). As a first step to define possible involvement of FGFs also in generation of ventral OPCs, we examined activation of the canonical MAPK pathway at the time of ventral OPC specification in chicken, i.e. between 5.5 and 6 days of development (E5.5/E6) (Soula et al., 2001; Zhou et al., 2001). We first analyzed expression of the active form of the signal-regulated protein kinase ERK1/2 (P-ERK1/2) on transverse spinal cord sections starting at E4 up to E7. At E4-E4.5, cells expressing P-ERK1/2 were detected in the ventral-most region of the ventricular zone (Fig. 1B). From E5, both intensity and dorso-ventral extent of the P-ERK1/2 immunostaining significantly increased (Fig. 1C, D), indicating temporal activation of the MAPK signaling pathway in ventral progenitor cells. Positioning of P-ERK1/2 positive cells with respect to Nkx2.2 showed that activation of ERK1/2 was initially restricted to Nkx2.2-expressing cells of the ventral-most p3 domain (Fig. 1E). At E5-E5.5, Nkx2.2 expression extended dorsally compared to earlier stages. P-ERK1/2 staining was then mostly detected in the dorsal-most Nkx2.2-expressing cells (Fig. 1F), indicating activation of the MAPK pathway in cells of the p* domain that has already being established at this stage (Zhou et al., 2001; Fu et al., 2002; Agius et al., 2004; Touahri et al., 2012). Similar patterns were observed at E6.5-E7 although the P-ERK1/2 signal extended more dorsally at these later stages (Fig. 1G).

Our data, showing activation of ERK1/2 in the ventral spinal cord at stages of OPC specification, indicate that, in addition to Shh, ventral progenitor cells perceived growth factor signals. Furthermore, the progression of ERK1/2 activation over time, starting in the p3 domain (prospective LFP cells) and further extending to dorsally-located progenitor cells (p* domain), was suggestive of a role of the MAPK signaling pathway both in LFP induction and OPC commitment.

### Ventral neural progenitors activate FGFR signaling at the time of OPC specification

We next sought to define whether ventral progenitor cells of the spinal cord are indeed targets for FGFs focusing on expression of FGFR and FGF target gene expression at stages of OPC specification. To date, four high affinity receptors for FGF (FGFRs1-4) have been identified. FGFRs 1-3 are known to be expressed in regions of the ventral embryonic forebrain that give rise to OPCs (Bansal et al., 2003). In the spinal cord, solely FGFR3 expression has been studied and it is preferentially expressed in progenitor domains dedicated to generate astrocytes but not OPCs (Pringle et al., 2003; Agius et al., 2004). We thus sought to establish expression patterns of FGFR1, 2 and 4. From E4-E4.5 until E6, mRNAs encoding for the three receptors were all detected in ventral progenitor cells with no gross change in their expression patterns over time (Fig. 2A-F). While *fgfr1* was uniformly and broadly expressed in progenitor cells (Fig. 2A, D), distinct expression levels for *fgfr2* and *fgfr4* were observed along the ventral to dorsal axis of the progenitor zone. *Fgfr2* mRNA was detected at higher level in a restricted domain of the ventral region, in a position very reminiscent to the source of OPCs (Fig. 2E). To confirm this, we performed double detection experiments using antibodies directed against Olig2 and Nkx2.2. Our data showed that Olig2-expressing cells as well as the dorsal-most Nkx2.2 positive cells were indeed included in the domain of high *fgfr2* expression (Fig. 2G, I). *Fgfr4* was detected in the entire ventral progenitor zone at E4-E4.5. However, at E5.5-E6, expression of this gene decreased in the ventral-most cells. Comparison with Olig2 and Nkx2.2 indicated that *fgfr4* expression overlapped the Olig2 domain as well as the dorsal-most region of the Nkx2.2 domain but was down-regulated in the ventral-most Nkx2.2-expressing domain (Fig. 2H, J) populated at this stage by LFP cells (Al Oustah et al., 2014). Together, these data indicate that ventral progenitor cells of both the p3 and pMN/p* domains can sense FGF signals at developmental stages of OPC specification.

**Figure 2:**
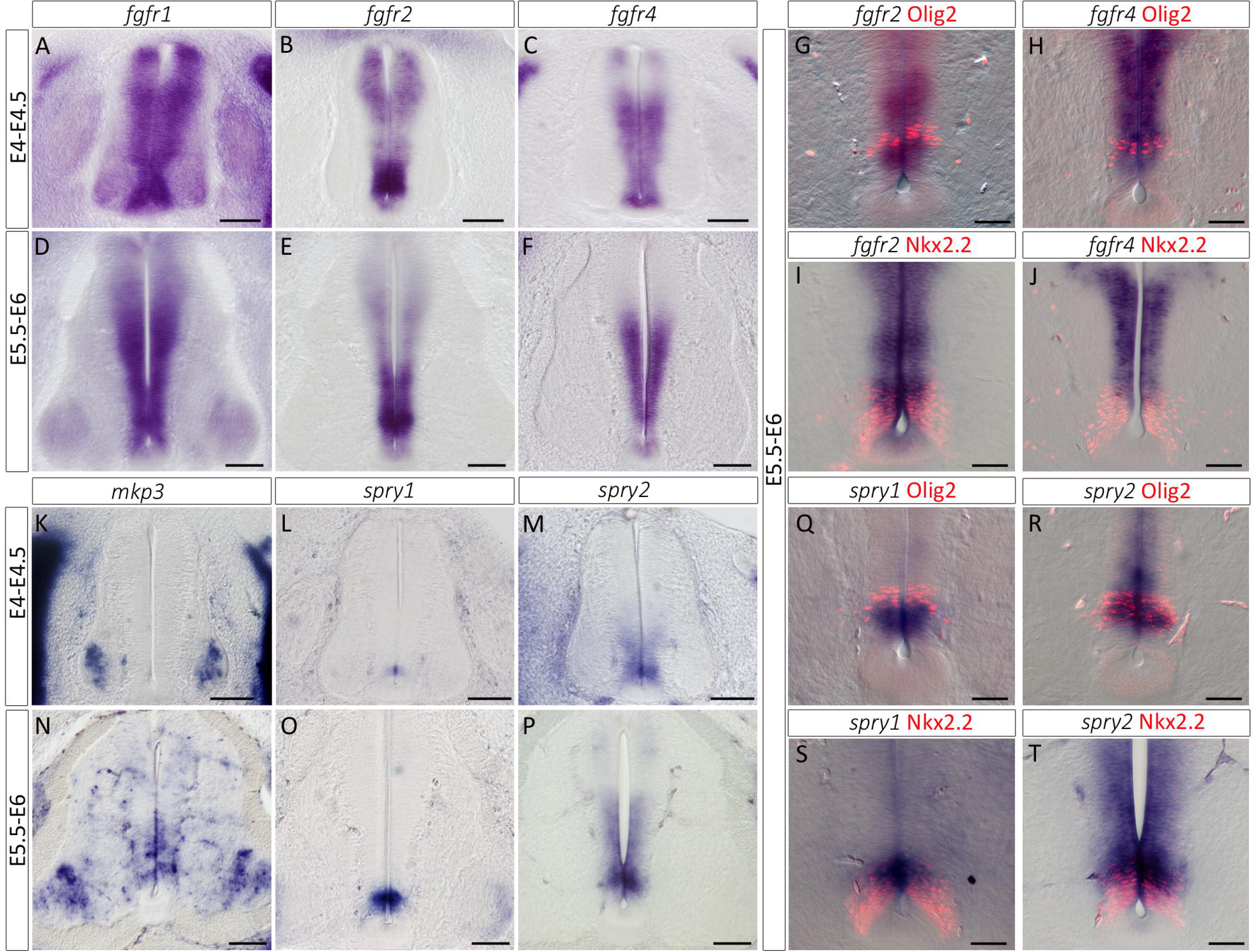
Expression of FGF receptors and FGF target genes at stages of ventral OPC specification in the spinal cord. **A-F**: Expression patterns of mRNA encoding for *fgfr1* (A, D), *fgfr2* (B, E) and *fgfr4* (C, F) on transverse spinal cord sections isolated before OPC specification (E4-E4.5) or at the onset of OPC generation (E5.5-E6). **G-H**: Double detection of Olig2 and *fgfr2* (G) or *fgfr4* (H) at E5.5-E6. Note expression of both mRNA in Olig2-positive progenitor cells at stages of OPC specification. **I-J**: Double detection of Nkx2.2 and *fgfr2* (I) or *fgfr4* (J) at E5.5-E6. Note that expression of *fgfr2* and *fgfr4* mRNAs is restricted to the dorsal-most region of the Nkx2.2 positive domain. **K-P**: Expression patterns of mRNA encoding for *mkp3* (K, N), *spry1* (L, O) and *spry2* (M, P) on transverse spinal cord sections showing that activation of *mkp3* in ventral progenitor cells occurs between E4-E4.5 and E5.5-E6 while expression of *spry1* and *spry2,* although detected at low level at E4-E4.5, is reinforced at E5.5-E6. **Q-R**: Double detection of Olig2 and *spry1* (Q) or *spry2* (R). **S-T**: Double detection of Nkx2.2 and *spry1* (S) or *spry2* (T) showing expression of both mRNA restricted to the dorsal-most Nkx2.2 positive progenitor cells at stages of OPC specification. Scale bars = 100μm in A-F and K-P, 50μm in G-J and K-T.

We then assessed activation of the FGFR signaling pathway by analyzing expression of the FGF target genes, *mkp3* (Eblaghie et al., 2003; Kawakami et al., 2003; Echevarria et al., 2005) and *sprouty1/2* (*spry1/2*) (Minowada et al., 1999; Kim and Bar-Sagi, 2004). Before OPC specification (E4/E4.5), we detected *mkp3* expression in pools of ventral neurons but not in progenitor cells (Fig. 2K). At this stage, low level of *spry1* expression was detected in a very ventral domain of the progenitor zone, adjacent to the MFP, while *spry2* mRNA extended in a broader domain of the ventral progenitor zone (Figure 2L, M). At E5.5-E6, expression of *mkp3* became apparent in cells of the ventral progenitor zone (Fig. 2N), indicating activation of FGFR signaling at stages of OPC specification. Accordingly, we detected higher level of *spry1/2* expression in ventral progenitor cells (Fig. 2O, P). Double detection of *spry1/2* and Olig2 or Nkx2.2 indicated preferential expression of these genes in Nkx2.2- and Olig2-expressing progenitor cells with higher levels of *spry1* and *spry2* expression in Nkx2.2 and Olig2 positive cells, respectively (Fig. 2 Q-T), suggesting domain specific responses to FGFR activation in the ventral progenitor zone.

Together, our data, showing that ventral progenitor cells of the p3 and p* domains both express FGFRs and up-regulate FGF target genes at stages of OPC specification, support the view that FGF signaling pathway might play a role in controlling generation of ventral spinal OPCs.

### Inactivation of FGFRs is sufficient to impair generation of ventral OPCs

To test the role of FGFs in ventral OPC specification, we turned to organotypic culture of chicken spinal cord explants that allows interfering with cell signaling precisely at the time of OPC induction (Agius et al., 2004; Danesin et al., 2006; Agius et al., 2010; Touahri et al., 2012). In this paradigm, brachial spinal cord explants are isolated prior to OPC specification (E4-E4.5) and plated in culture as an opened-book. With this system, the temporal fate of Olig2 progenitor cells is maintained and OPC migration in the mantle zone can be observed after two days in culture, a stage equivalent to E6-E6.5 *in vivo* (Agius et al., 2004; Danesin et al., 2006; Touahri et al., 2012). To address the requirement of FGF activity for ventral OPC generation, we inactivated FGF receptors (FGFRs) prior to OPC commitment (E4-E4.5) and analyzed OPC production after two days in culture. FGFR inactivation was performed either incubating explants with the FGFR inhibitors SU5402 and PD166866 (Mohammadi et al., 1997; Panek et al., 1998) or by overexpressing of a dominant-negative form of FGFRs (dnFGFR) that completely lacks the intracellular tyrosine kinase domain (Amaya et al., 1991) using the electroporation method. As a first step, we controlled efficient inactivation of FGFRs. SU5402 or PD166866 were added to the culture medium at the time of plating and expression of P-ERK1/2 and *spry1/2* were analyzed two days later. In control explants, P-ERK1/2 and *spry1/2* expressions were detected in ventral progenitor cells (Fig. 3A, C, E), identifying cells with patterns very similar to those observed on transverse spinal cord sections at E6. After treatment with FGFR inhibitors, we found that both P-ERK1/2 immunostaining and *spry1/2* expression were nearly abolished (Fig. 3B, D, F). Similarly, electroporation of the dnFGFR expression vector, performed just prior to spinal cord explantation, prevented both ERK1/2 activation and *spry1/2* expression while electroporation of a control vector had no effect (Fig. 3G-J). These results, beyond validating the approach, support the view that FGFRs are indeed responsible for stimulation of MAPK activity and *spry1/2* expression in the ventral spinal cord.

**Figure 3:**
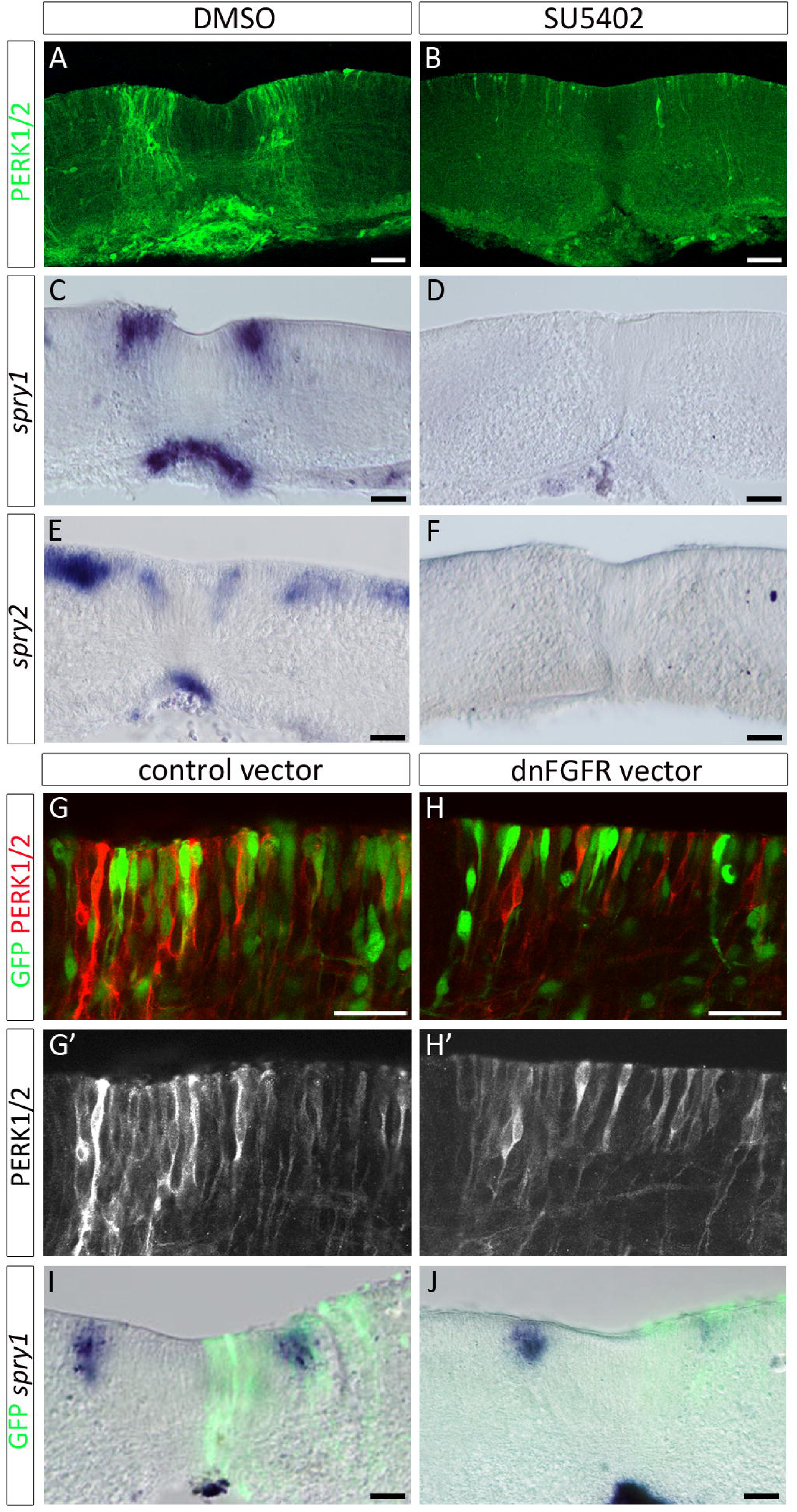
Inactivation of FGFRs results in defective activation of ERK1/2 and in down-regulation of *spry1/2* expression in spinal cord explants. **A-B**: Immunodetection of P-ERK1/2 on transverse sections of spinal cord explants isolated at E4.5 and cultivated as opened-book explants (progenitor cell layer on top) for 2 days in control condition (A) or in presence of SU5402 (B). **C-F**: Detection of *spry1* (C, D) and *spry2* (E,F) in control condition (C, E) or in presence of SU5402 (D, F). **G-H**: Higher magnification of the ventral progenitor zone (floor plate cells are on the left) showing immunodetection of PERK1/2 (red) after electroporation of the control vector (green in G) or the dnFGFR vector (green in H). **I-J**: Detection of *spry1* mRNA after electroporation of the control vector (green in I) or the dnFGFR vector (green in J). Scale bars=50μm.

We next examined OPC generation after FGFR inactivation. For this, we used the O4 antibody, a specific and early marker of OPCs in chicken (Ono et al., 1995; Miller et al., 1997; Soula et al., 2001) and Olig2, whose expression in the mantle zone allows identifying OPCs (Lu et al., 2000; Takebayashi et al., 2000; Zhou et al., 2000). In control explants, O4 positive and Olig2-expressing OPCs were detected both in the progenitor zone and mantle layer, indicating ongoing generation of OPCs (Fig. 4A). By contrast, only few O4 positive and Olig2-expressing cells were detected in the mantle zone of explants incubated with SU5402 or PD166866 (Fig. 4B). Cell counting indicated that the number of OPCs were reduced by more than 80% in presence of inhibitors compared to control explants (Fig. 4E, F), indicating that inhibition of FGFR activity is sufficient to prevent generation of ventral OPCs. Similarly, overexpression of the dnFGFR led to a greater than 5-fold reduction in the number of OPCs in the electroporated side of the explants compared to the non-electroporated side or to explants electroporated with a control vector (Fig. 4C-D, G-H). Importantly, cell proliferation studies showed that overall BrdU incorporation in the presence of SU5402 or PD166866 was not significantly different from controls (Fig. 5C-E). Similarly, cell survival, assessed by detection of the activated form of Caspase 3, was not affected by treatment with the inhibitors (Fig. 5A, B).

**Figure 4:**
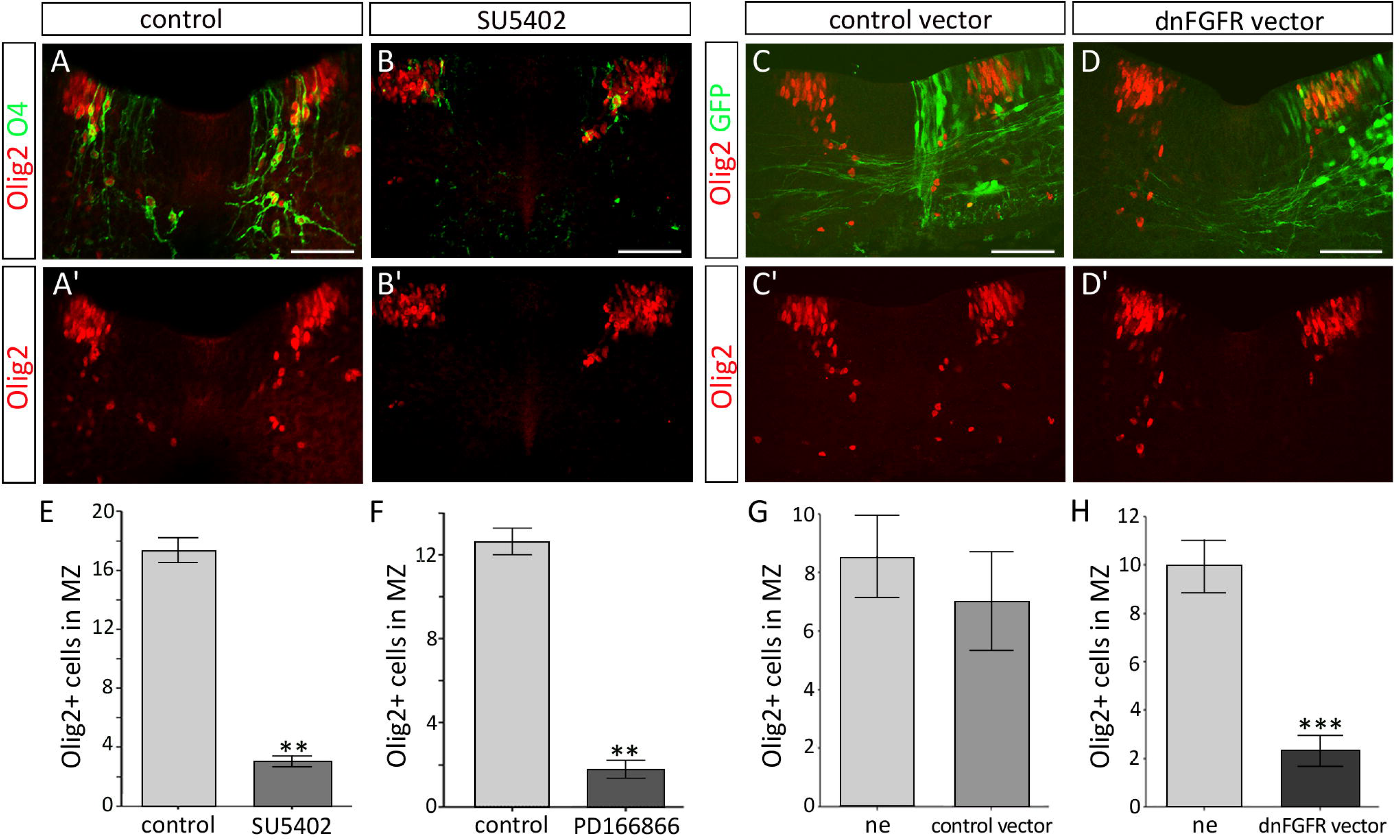
Inactivation of FGFRs impairs OPC generation. **A-B**: Immunodetection of O4 (green) and Olig2 (red) on transverse sections of spinal cord explants cultivated for 2 days in control condition (A) or in presence of SU5402 (B). Note that while O4/Olig2-positive OPCs have already invaded the mantle zone in control explants, very few develop in presence of the inhibitor. **C-D**: Immunodetection of Olig2 (red) after electroporation of control (green in C) or dnFGFR (green in D) vectors. Note that very few Olig2-positive cells have migrated in the mantle zone on the side of the explant electroporated with the dnFGFR vector. **E-F**: Quantification of Olig2-positive cells in the mantle zone in control conditions (n=7 for each essay) or in presence of the FGFR inhibitors SU5402 (n=8, E) or PD166866 (n=6, F). **G-H**: Quantification of Olig2-positive cells migrating in the mantle zone on each side of the explants after electroporation of control (n=4, G) or dnFGFR (n=9, H) vectors. Results are presented as mean number of cells ± sem (**p≤0.01; *** p≤0.001). ne: non-electroporated side of explants. Scale bars = 50μm.

**Figure 5:**
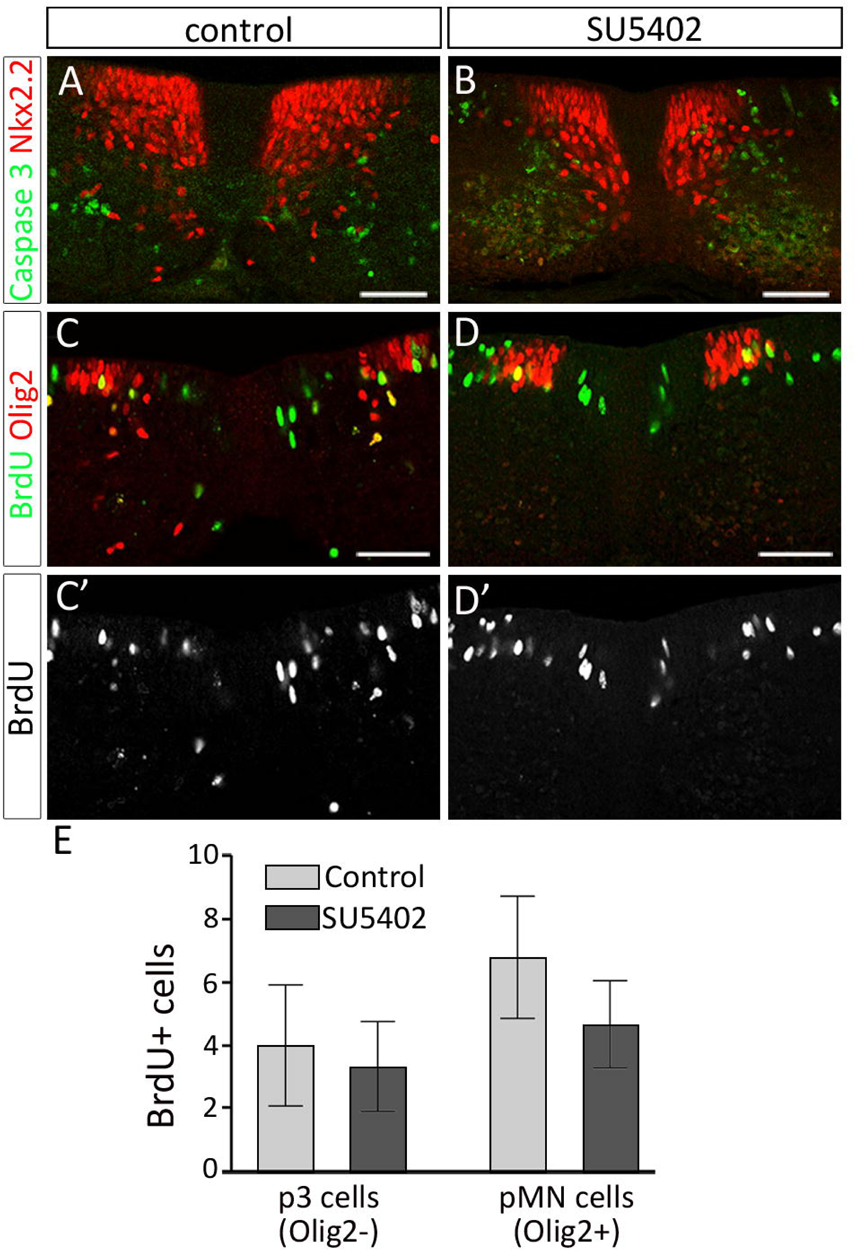
Inactivation of FGFRs does not affect cell survival and proliferation of spinal cord cells. **A-B**: Detection of clived caspase 3 and Nkx2.2 on transverse sections of spinal cord explants cultivated in control condition (A) or in presence of SU5402 (B). **C-D**: Detection of BrdU (green) and Olig2 (red) in control condition (C) or in presence of SU5402 (D). BrdU pulse has been performed at 32 hours after plating. **E**: Quantification of BrdU-positive cells in the p3 (Olig2-negative) and pMN (Olig2-positive) domains of the progenitor zone on explants cultivated in control condition or in presence of SU5402. Scale bars = 50 μm.

Together, our data, showing that inactivation of FGFRs is sufficient to inhibit OPC development, indicate that FGFR signaling is required for proper generation of OPCs in the ventral spinal cord and that this function does not rely on the control of cell proliferation or cell survival.

### FGFR activity is required for OPC specification in the ventral spinal cord

We next sought to define whether FGFRs contribute to commit Olig2-expressing progenitor cells to the OPC fate. To test this, we performed double immunostainings of Olig2 and Nkx2.2, whose coexpression in progenitor cells of the ventral spinal cord is a hallmark of OPC specification (Zhou et al., 2001; Fu et al., 2002; Agius et al., 2004; Danesin et al., 2006; Touahri et al., 2012; Al Oustah et al., 2014), on explants treated with FGFR inhibitors or explants overexpressing dnFGFR. After two days in culture, Nkx2.2/Olig2-coexpressing cells were detected in the progenitor zone forming the p* domain but also in newly produced OPCs that have emigrated in the mantle zone (Fig. 6A). As in the previous experiments, very few Nkx2.2/Olig2-coexpressing OPCs were detected in the mantle zone of explants treated with SU5402 (Fig. 6B). Examination of the progenitor zone showed that, while position and size of the Olig2 expression domain were not affected, the dorsal extent of the Nkx2.2-expressing domain was reduced compared to control explants (Fig. 6C-D). Cell counting showed a significant decrease in the number of Olig2/Nkx2.2-coexpressing progenitor cells in explants treated either with SU5402 or PD166866 compared to control explants (Fig. 6E), indicating defective generation of the p* domain in the presence of FGFR inhibitors. Similar results were obtained when FGFR inhibition was performed using dnFGFR overexpression in ventral progenitor cells (Fig. 6F, G).

**Figure 6:**
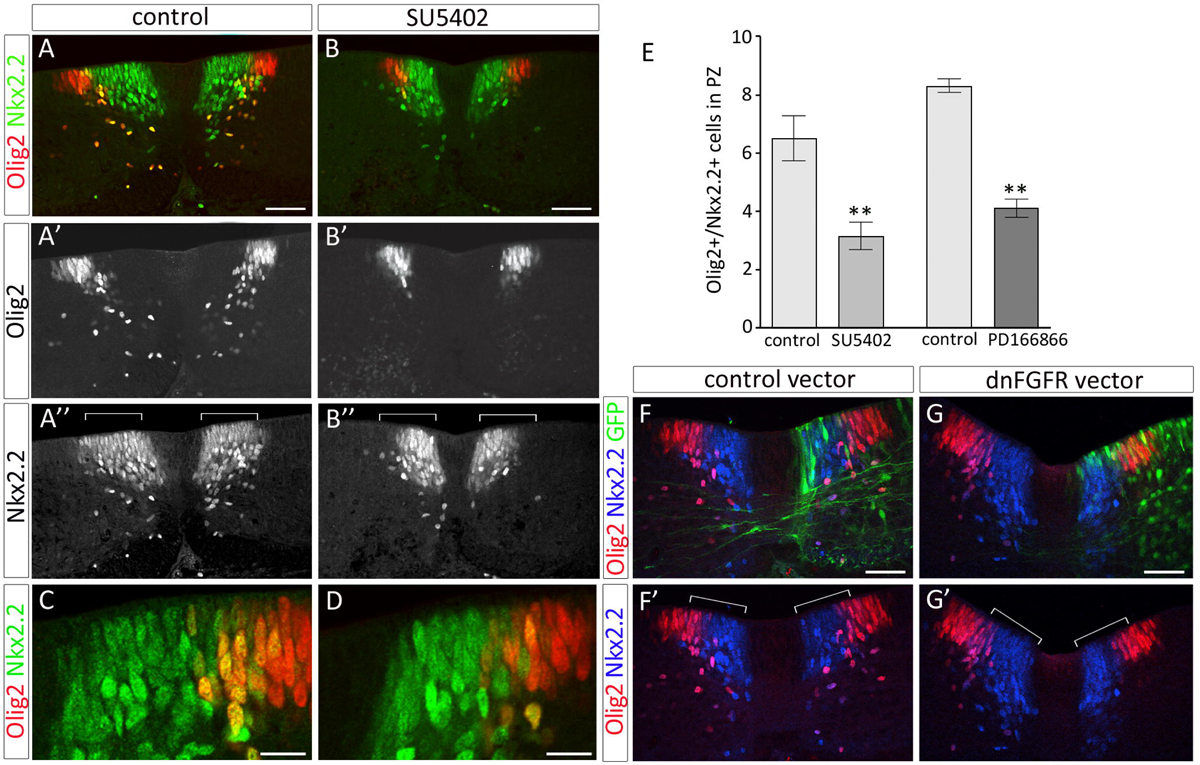
Inactivation of FGFRs impairs OPC specification. **A-B**: Immunodetection of Olig2 (red) and Nkx2.2 (green) on transverse sections of spinal cord explants cultivated for 2 days in control condition (A) or in presence of SU5402 (B). Note reduced dorsal extent of the Nkx2.2-positive domain (brackets) in SU5402 treated explant (B’’) compared to control explant (A’’). **C-D**: Higher magnification of the progenitor zone showing reduced number of Olig2/Nkx2.2-positive cells in the progenitor zone of explants treated with SU5402 (D) compared to control explants (C). **E**: Quantification of NKx2.2/Olig2-positive cells in the progenitor zone (PZ) of explants cultivated in control conditions (n=7 for each essay) or in presence of SU5402 (n=8) or PD166866 (n=6). Results are presented as mean number of cells ± sem (**p≤0.01). **F-G**: Immunodetection of Olig2 (red) and Nkx2.2 (blue) after electroporation of control (green in F) or dnFGFR (green in G) vectors. Note reduction of the dorsal extent of the Nkx2.2 positive domain in the side of explant electroporated with the dnFGFR vector compared to the non electroporated side of the explant or with explant electroporated with the control vector (brackets). Note also that migrating Olig2/Nkx2.2 double-labeled cells are not detected in the side of explant electroporated with the dnFGFR vector (G’). Scale bars = 50 μm in A, B, F, G and 25 μm in C, D.

These data, showing that inhibition of FGFR signaling activity is sufficient to prevent formation of the p* domain, highlight the key role of FGFRs in controlling specification of OPCs in the ventral spinal cord.

### Formation of the lateral floor plate requires activation of FGFR signaling

We know from our previous work that formation of the LFP by up-regulation of *shh* in Nkx2.2-expressing cells of the p3 domain is temporally correlated with time commitment of Olig2 progenitor cells to the OPC fate (Al Oustah et al., 2014; Danesin and Soula, 2017). However, whether LFP formation contributes to create the burst of Shh signaling activity required for OPC specification remains elusive. Based on our present data, showing that Nkx2.2-expressing cells of the p3 domain express FGFRs and activate P-ERK1/2 prior to LFP induction (E4/E4.5, Figs. 2 and 3), we hypothesized that FGFs might contribute to induce OPCs indirectly, by stimulating Shh expression within prospective LFP cells. To test this, we analyzed the effects of FGFR inactivation on Shh expression using the spinal cord explant paradigm. As a first step, we asked whether LFP forms properly in explanted spinal cords. At E4, corresponding to the stage of spinal cord explantation, *shh* mRNA expression was restricted to MFP cells (Fig. 7A). After 9 hours in culture, the time required for explant attachment to the substratum, detection of *shh* mRNA was still limited to ventral medial cells of the MFP (Fig. 7B). By contrast, 1 day after plating, equivalent to E5-E5.5 *in vivo, shh* expression extended in dorsal direction, i.e. laterally in opened-book explants (Fig. 7C). Comparison with Nkx2.2 expression at an equivalent culture time indicated that *shh* expression encompassed the Nkx2.2 positive p3 domain (Fig. 7C, E). After 2 days in vitro, *shh* mRNA was still detected in LFP cells and, expectedly, the domain of *shh* expression did not span the dorsal-most part of Nkx2.2 positive domain corresponding, at that time, to the p* domain (Fig. 7D, F). Therefore, LFP develops properly in spinal cord explants and its induction occurs within the first day of culture, in agreement with its period of formation *in vivo* (Al Oustah et al., 2014). Strikingly, SU5402 treatment, while not modifying *shh* expression in MFP cells, nearly abolished extension of *shh* expression in cells of the p3/LFP domain at either 1 or 2 days in culture (Fig. 7G, H). Confirming these data, overexpression of the dnFGFR in cells of the p3 domain appeared sufficient to prevent activation of *shh* expression in these cells (Fig. 8A). We next limited SU5402 treatment to the first day of culture, i.e. prior to LFP induction, or to the second day of culture, after LFP formation, and analyzed, in either case, *shh* expression at day 2. We found that inhibition of FGFR was sufficient to prevent up-regulation of *shh* when performed prior to LFP induction (Fig. 7I) but had no influence of *shh* expression after the time of LFP induction (Fig. 7J). Therefore, FGFR activation is required for induction but not for maintenance of LFP cells. We next asked whether LFP induction, as reported for induction of the MFP (Sasai et al., 2014), resulted from concerted activities of FGFs and Shh which is still secreted by MFP cells at the time of LFP formation. We then incubated spinal cord explants with cyclopamine, a potent inhibitor of the Shh signaling pathway (Cooper et al., 1998; Incardona et al., 1998) and analyzed LFP formation. As previously reported (Agius et al., 2004), after 2 days in culture, cyclopamine treatment inhibited expression of Olig2, known to require Shh activity to be maintained, but also prevented dorsal extension of Nkx2.2 expression (Fig. 7K, L). By contrast, this treatment had no effect on *shh* expression, indicating that LFP forms properly in cyclopamine treated explants (Fig. 7 M, N). Therefore, induction of the LFP is independent of Shh but depends on FGFR signaling. These data are in agreement with our data showing that cells of the p3 domain express activated forms of ERK1/2 as well as FGF target genes (see Figs 1 and 2) and down-regulate the Shh receptor Patched over time (Touahri et al., 2012). We next took advantage of this paradigm to assess the requirement of LFP formation for OPC induction. For this, we inactivated FGFRs specifically in prospective LFP cells by targeting electroporation of the dnFGFR vector to the p3 domain. We then analyzed OPC generation in these explants using Olig2 and Nkx2.2 double immunostaining. We found that inactivation of FGFR signaling specifically in cells of the p3 domain was sufficient to prevent OPC generation (Fig. 8B-D). Of note, inactivation of FGFRs specifically in the p3 domain prevented dorsal extension of the Nkx2.2-expressing domain but did not impair Olig2 expression that was still detected in cells of the adjacent pMN domain (Fig. 8C). These data demonstrate that establishment of the LFP is required to trigger OPC generation and indicate that the prime function of LFP cells is to activate Nkx2.2 expression to form the p* domain but not to maintain Olig2 expression in pMN progenitor cells, the latter function being probably assumed by Shh still provided by MFP cells in this experimental context.

**Figure 7:**
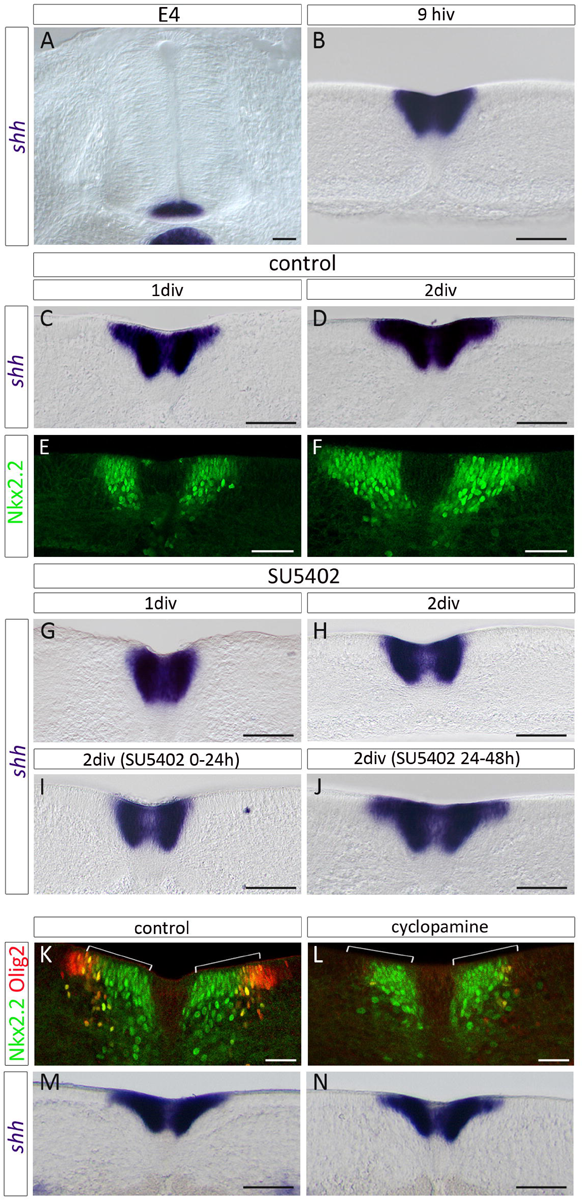
LFP induction requires FGFR activation. **A**: Expression pattern of *shh* mRNA on transverse section of E4 brachial spinal cord showing *shh* expression restricted to MFP cells at this stage. **B-D**: Time course of *shh* expression in spinal cord explants dissected at E4-4.5 and cultivated for 9 hours (B), 1 day (C) or 2 days (D). **E-F**: Immunodetection of Nkx2.2 on transverse sections of explants cultivated for 1 day (E) or 2 days (F). Note that dorsal extension of *shh* expression overlaps the Nkx2.2-positive domain to form the LFP at day 1 (compare C and E) while dorsal extension of the Nkx2.2 domain to form the p* domain is detected only at day 2 (F). **G-H**: Time course of *shh* expression through spinal cord explants cultivated in presence of SU5402 for 1 day (G) or 2 days (H-J). Note that dorsal extension of *shh* expression to form the LFP fails to occur in presence of the FGFR inhibitor. **I-J**: Expression of *shh* in spinal cord explants cultivated for 2 days and treated for a time period of 24 hours by SU5402, either during the first (I) or the second day in culture (J). Note that LFP fails to form even when the treatment is interrupted after the first day in culture (I), whereas it forms readily when the inhibitor is applied only during the second day of culture (J). **K-N**: Transverse section of control explants (K, M) or explants treated with cyclopamine for 2 days (L, N) and immunolabeled for Olig2 and Nkx2.2 (K, L) or processed for *shh* mRNA detection (M, N). Note that cyclopamine treatment inhibits Olig2 expression and prevents dorsal extension of the Nkx2.2 domain (brackets, L), whereas the treatment has no effect on dorsal extension of *shh* mRNA expression (N). hiv = hour in vitro, div = day in vitro. Scale bars = 50 μm.

**Figure 8:**
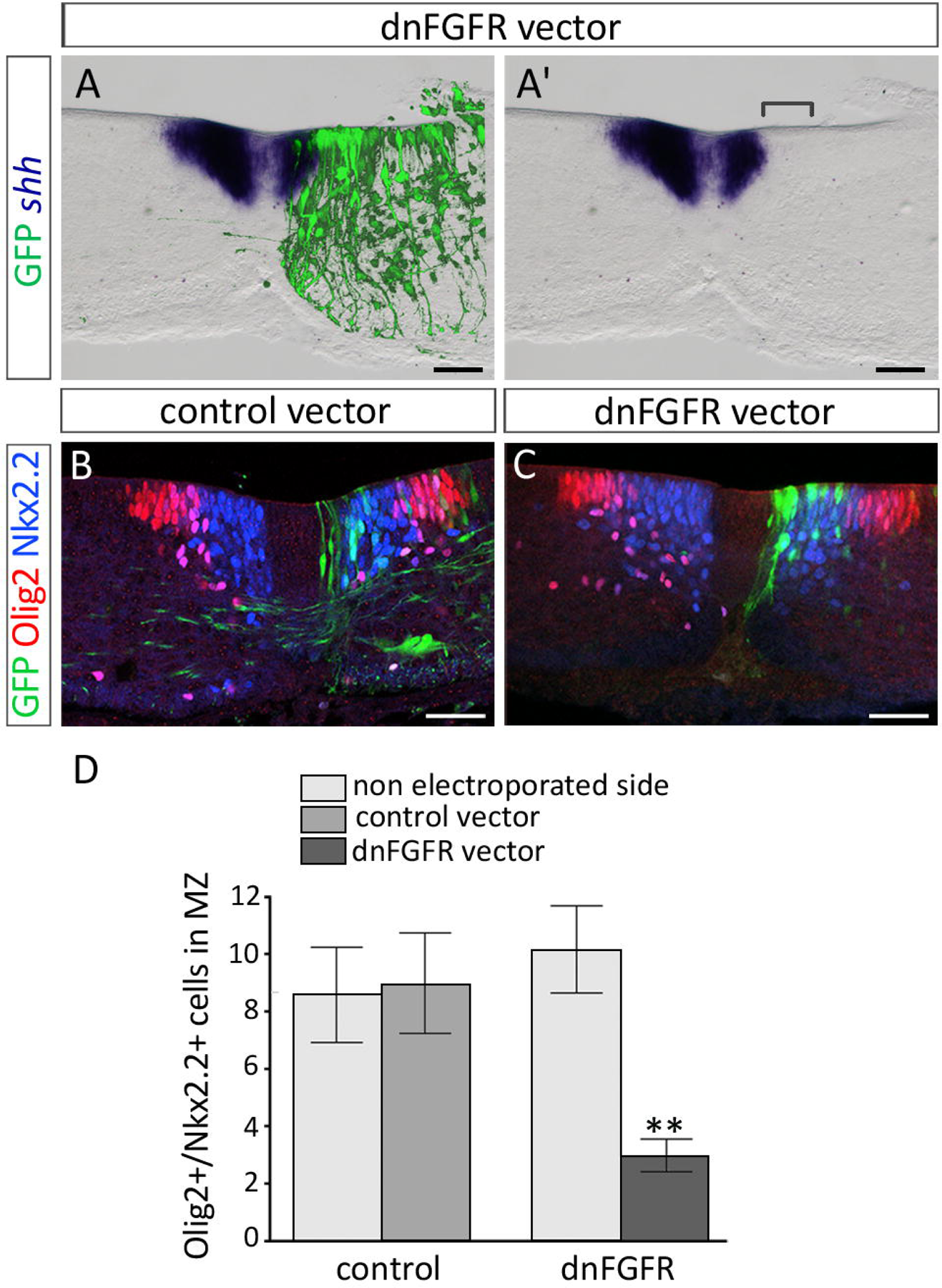
Cell autonomous activation of FGFRs on p3 progenitor cells is required for activation of Shh expression. **A**: Relative location of *shh* mRNA-expressing cells (purple) and dnFGFR electroporated cells (green) on transverse sections of spinal cord explant cultivated for 2 days. Note inhibition of *shh* expression in the ventral-most dnFGFR-overexpressing cells (bracket in A’). **B-C**: Double immunodetection of Olig2 (red) and Nkx2.2 (blue) in explants electroporated with the control vector (green in B) or with the dnFGFR vector (green in C). Note that limiting overexpression of dnFGFR to the Nkx2.2-positive domain (C) is sufficient to prevent generation of Olig2/Nkx2.2-coexpressing OPCs. **D**: Quantification of Olig2/Nkx2.2-positive cells in the marginal zone (MZ) of non-electroporated sides of explants and sides of explants electroporated with the control (n=4) or dnFGFR (n=5) vectors. Results are presented as mean number of cells ± sem (**p≤0.01). Scale bars = 50 μm.

Together, our data highlight a key and indirect function for FGFR signaling in triggering ventral OPC specification, through the induction of a secondary source of Shh required for formation of the p* domain.

### FGFR activation on Olig2 progenitors is required for OPC generation

As Olig2 progenitor cells of the pMN domain also express FGFRs and activate expression of FGF target genes as they start generating OPCs, we hypothesized that FGFRs might also control OPC commitment, acting cell-autonomously on these cells. To test this, we selected explants in which Olig2 progenitor cells, but not the more ventrally located Nkx2.2-expressing ones, had been targeted by electroporation of the dnFGFR expression vector. Accordingly, after 2 days in culture, we observed equivalent ventral to dorsal extension of Nkx2.2 expression domain either in dnFGFR electroporated side of explants, non electroporated explants or in explants electroporated with the control vector (Fig. 9A, B). Despite this, overexpression of dnFGFR in Olig2 progenitor cells reduced the population of Nkx2.2./Olig2-coexpressing OPCs emigrating in the mantle layer (Fig. 9B). Noticeably, in the dnFGFR electroporated side of these explants, we observed a marked reduction in the expression of Olig2 while Nkx2.2 expression was unaffected. More specifically, the number of cells expressing high level of Olig2 appeared reduced when the dnFGFR expression vector, instead of the control vector, was used (Fig. 9C), suggesting that FGFR activity on Olig2-expressing cells contribute to maintain high level of Olig2 expression at stages of OPC commitment, a necessary condition for triggering progenitor cells to follow an OPC fate (Novitch et al., 2001; Qi et al., 2003; Liu et al., 2007).

**Figure 9:**
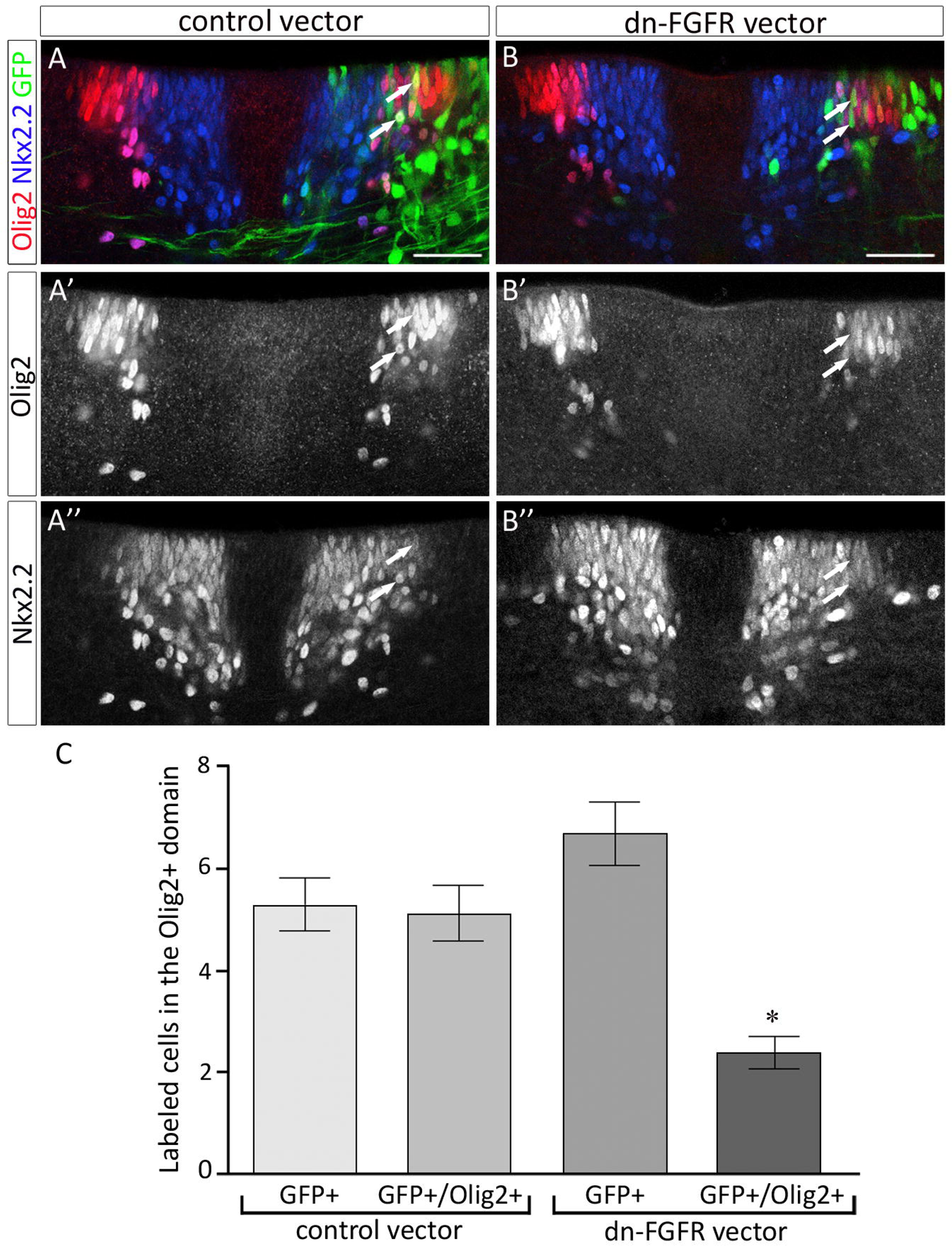
Cell autonomous activation of FGFRs on pMN progenitor cells is required to maintain Olig2 expression at the time of OPC specification. **A-B**: Immunodetection of Olig2 (red) and Nkx2.2 (blue) on transverse sections of explants electroporated with the control vector (green in A) or with the dnFGFR vector (green in B) and cultivated for 2 days. Note the reduced number of Olig2/Nkx2.2-positive cells generated in the dnFGFR electroporated side of explant in B. Note also diminished level of Olig2 expression in progenitor cells electroporated with the dnFGFR vector (arrows in B) compared to those electroporated with the control vector (arrows in A). **C**: Quantification of GFP-positive and GFP/Olig2-positive cells in the Olig2 progenitor domain in control (n=3) and dnFGFR electroporated (n=4) explants. Results are presented as mean number of cells ± sem (* p≤0.03). Scale bars = 50 μm.

Together, these data, showing that inactivation of FGFR specifically on Olig2 progenitors is sufficient to impair OPC production, highlighted a cell autonomous role of FGFR on Olig2-expressing progenitor cells and point to a role of FGFRs in maintaining Olig2 expression.

### FGF8 and Shh work together on neural progenitor cells to promote co-expression of Olig2 and Nkx2.2

Olig2 expression is known to depend on low doses of Shh for its induction (Lu et al., 2000; Zhou et al., 2000). It is also known to be repressed by high levels of Shh signaling at early stages of neural tube patterning, a process mediated by the activation of Nkx2.2 (Novitch et al., 2001; Jeong and McMahon, 2005; Stamataki et al., 2005; Lei et al., 2006; Dessaud et al., 2007; Lek et al., 2010). As mentioned above, the situation is quite different at stages of p* domain formation since up-regulation of Nkx2.2 no more represses Olig2 expression. The lowered intensity of the Olig2 signal observed in dnFGFR overexpressing pMN cells at the time of OPC specification therefore led us to consider the possibility that FGFR activation might be a key determinant for providing a cell context allowing maintenance of Olig2 expression as Shh signaling rises in the ventral spinal cord. To test whether FGFs might influence Olig2 expression in a context of high Shh signaling activity, we turned to the early neural tube and monitored expression of Olig2 and Nkx2.2 after electroporation of constructs, used alone or in combination, that encode for Shh or for FGF8, a ligand known to bind and activate FGFR1-4 in vertebrates (Mason, 2007). When the FGF8 expression vector was electroporated alone, expression of Nkx2.2 or Olig2 was not obviously altered and we did not detect ectopic expression of either protein in the dorsal neural tube (Fig. 10A, D). As expected, misexpression of Shh alone induced ectopic expression of Nkx2.2 and Olig2 in progenitor cells of the dorsal neural tube (Fig. 10B, E). Most of these ectopic cells expressed either Olig2 or Nkx2.2, likely reflecting cell exposure to distinct levels of Shh, and only very few of them co-expressed Nkx2.2 and Olig2 (Fig. 10E). By contrast, when both FGF8 and Shh were misexpressed in the neural tube, most if not all induced cells co-expressed Olig2 and Nkx2.2 (Fig. 10C, F). Thus, joint exposure of progenitor cells to FGF8 and high levels of Shh promotes induction of Olig2 and Nkx2.2 coexpressing cells, supporting the idea that FGFR activation contributes to maintain Olig2 expression in progenitor cells as they activate Nkx2.2 in response to high threshold Shh signaling.

**Figure 10:**
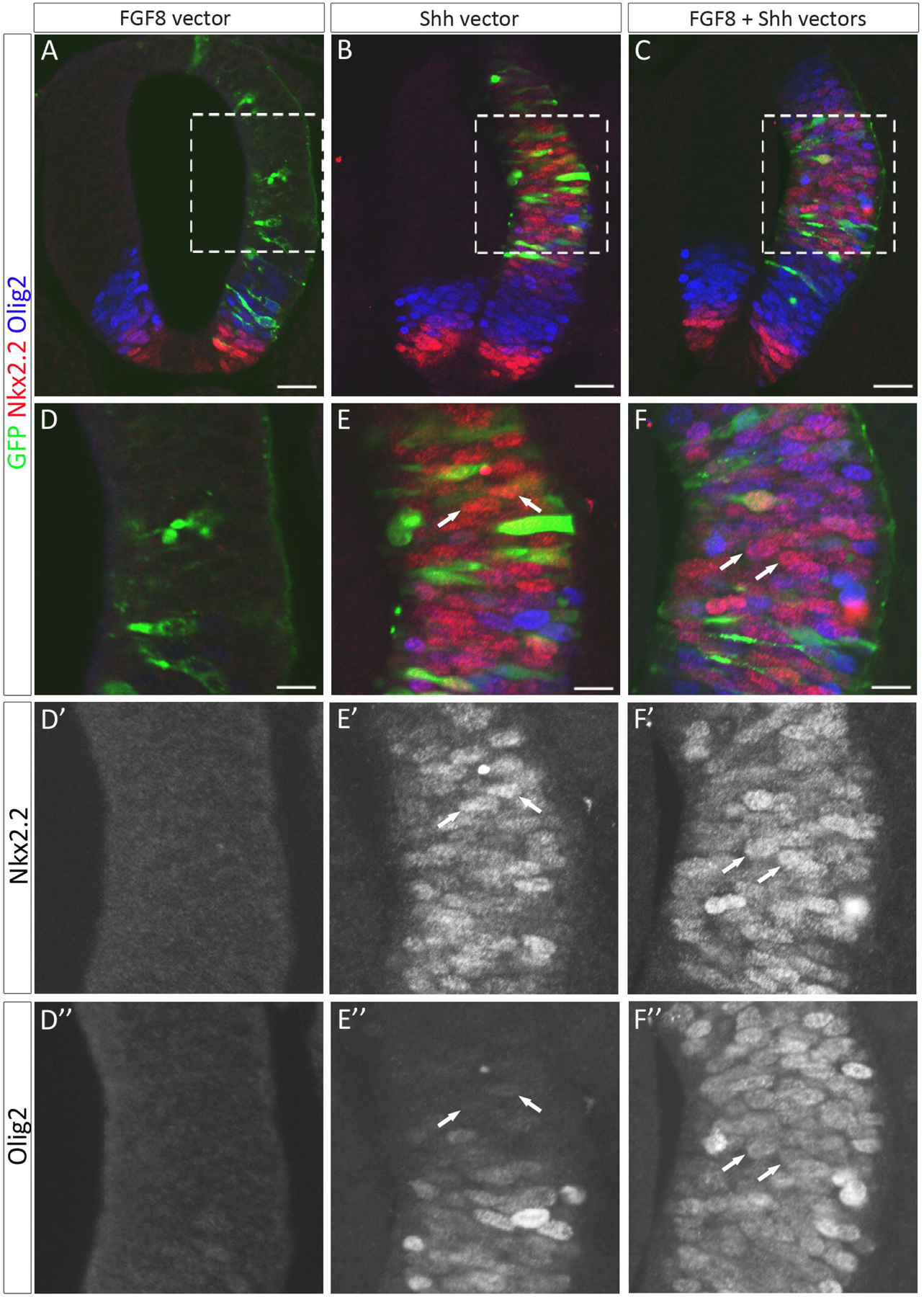
Coexposure of neural progenitor cells to FGF8 and Shh signals favors co-expression of Olig2 and Nkx2.2. **A-F**: Immunodetection of Olig2 (blue) and Nkx2.2 (red) on transverse E2.5 neural tube sections after electroporation at E1.5 with FGF8 (A, D), Shh (B, E) or both FGF8 and Shh expression vectors (C, F). D, E and F show higher magnification of the areas framed in A, B and C, respectively. Note that Olig2 expression is detected in most of the Nkx2.2 positive cells induced by overexpression of both FGF8 and Shh (arrows in F, F’ and F’’) but not in Nkx2.2 positive cell induced by overexpression of Shh alone (arrows in E, E’ and E’’). Scale bars = 50 μm in A-C, 25 μm in D-F.

We then postulated that, according to this conclusion, overexpression of FGF8 might be sufficient to promote Olig2/Nkx2.2 coexpression in ventral progenitor cells at the time of LFP formation, i.e. when Shh signaling activity rises in the ventral spinal cord. To test this, we overexpressed FGF8 at E4-E4.5 and analyzed expression of Olig2 and Nkx2.2 after 2 days in culture to cover the time period of Shh signaling activation. Contrasting with data got in the earlier neural tube, we found that electroporation of the FGF8 expression vector in these later stages, was sufficient to promote induction of Olig2 and Nkx2.2 coexpression in ventral progenitor cells compared to explants electroporated with control vector (Fig. 11A, B). To define whether the inductive activity of FGF8 at these later stages depends of Shh, we performed similar experiments incubating explants with cyclopamine. In agreement with previous report (Agius et al., 2004), we found that, after 2 days in culture in explants electroporated with the control vector, cyclopamine treatment down-regulated Olig2 expression in pMN progenitor cells and prevented dorsal extension of the Nkx2.2 domain thus totally abolishing OPC production (Fig. 11C). Similar experiments were then performed in explants electroporated with the FGF8 vector. In this context, the Olig2 progenitor domain as such was still not detectable either in FGF8 electroporated or non electroporated sides of explants and very few Olig2/Nkx2.2 coexpressing cells were found emigrating in the mantle layer in the electroporated side of explants (Fig. 11D). Therefore, FGF8 overexpression, although able to rescue generation of very few Nkx2.2/Olig2-coexpressing cells, was not sufficient to prevent cyclopamine-dependent down-regulation of Olig2 in progenitor cells.

**Figure 11:**
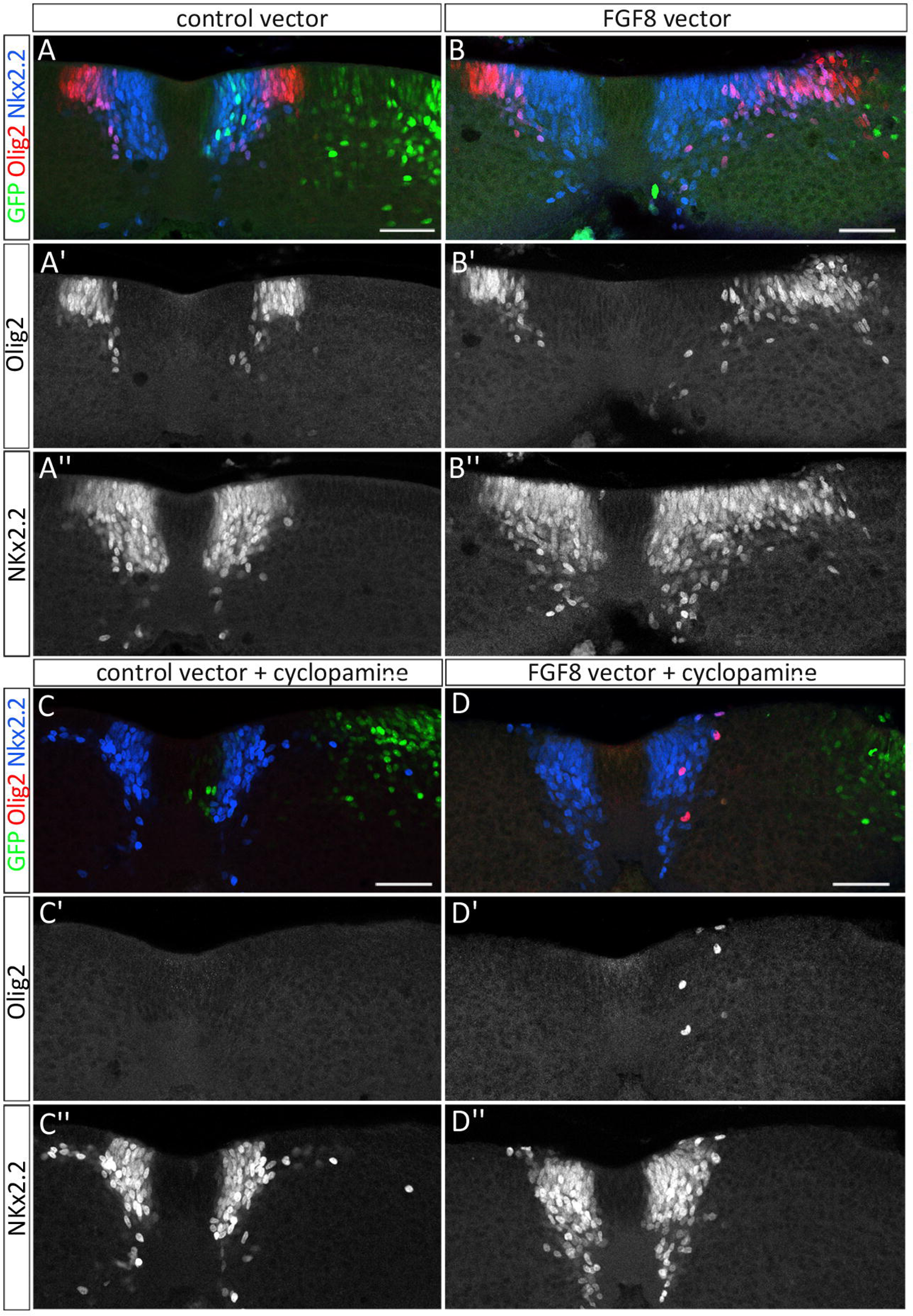
FGF8 promotes OPC specification following a Shh-dependent process. **A-B**: Immunodetection of Olig2 (red) and Nkx2.2 (blue) on transverse sections of spinal cord explants cultivated for 2 days after electroporation of the control vector (green in A) or the FGF8 vector (green in B) at E4-E4.5. **C-D**: Immunodetection of Olig2 (red) and Nkx2.2 (blue) on transverse sections of spinal cord explants electroporated with the control vector (green in C) or the FGF8 vector (green in D) and cultivated for 2 days in presence of cyclopamine. Scale bars = 50 μm.

Together, these data provide additional support to the view that commitment of ventral spinal OPCs depends on combined activation of FGF and Shh signaling, a cell context favoring Olig2 and Nkx2.2 coexpression.

### FGF8 is a good candidate for triggering induction of ventral OPCs *in vivo*

Our results raise the question of which cell subtype delivers FGF(s) signal(s) to initiate OPC commitment. FGF8 being efficient in its ability to induce OPCs, we analyzed its expression over spinal cord development. At E2.5, during ongoing generation of neurons, we did not find *fgf8* expression in the spinal cord (Fig. 12A). By contrast, from E4, high levels of *fgf8* expression were detected in sub-set of neurons located at the lateral edge of the progenitor zone. These neurons were at first restricted to intermediate regions of the developing spinal cord but they further shifted ventrally to position themselves in the vicinity of the p3 and pMN domains (Fig. 12B, C). Comparison with the expression pattern of Pax2, a specific marker of interneurons in the ventral spinal cord (Burrill et al., 1997), was supportive of *fgf8* being expressed in ventral interneuron sub-types (Fig. 12D-F). Activation of FGF8 in a subset of ventral neurons at the right time and place makes this ligand a good candidate to cooperate with Shh for OPC induction in the ventral spinal cord. Unexpectedly, these data also open the possibility that ventral interneurons might be players of Olig2 progenitor decision to engage in OPC generation.

**Figure 12:**
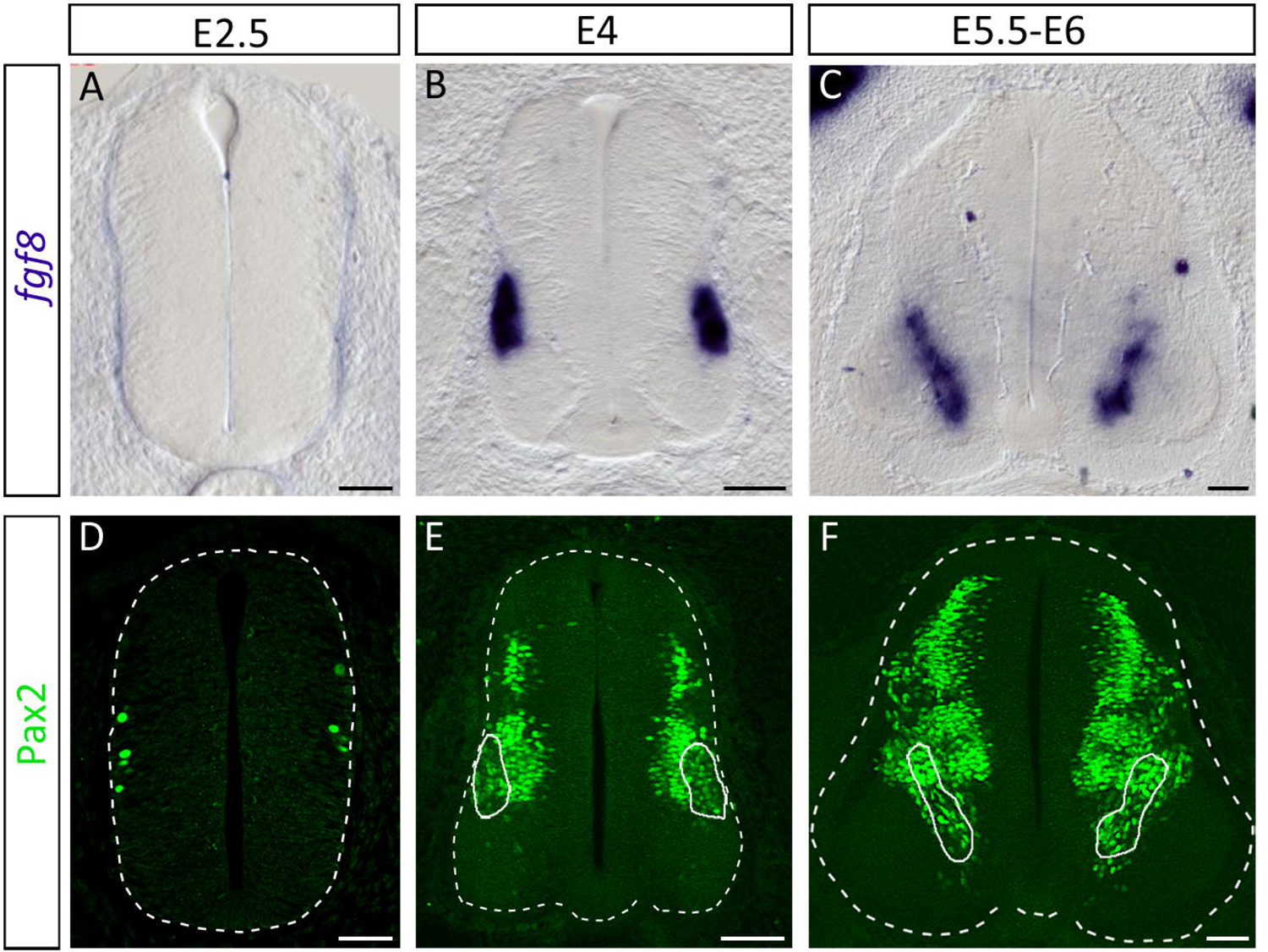
FGF8 expression is activated in ventral neurons at the time of OPC specification. **A-C**: Time course of *fgf8* expression on transverse sections of brachial spinal cords isolated at E2.5 (A), E4 (B) and E5.5-E6 (C). **D-F**: Immunodetection of Pax2 depicting spinal cord interneurons at E2.5 (D), E4 (C) and E5.5-E6 (D). Note that *fgf8* expressing cells correspond to a subpopulation of Pax2-positive cells (circled in E, F). Scale bars = 50 μm in A, D and 100 μm in B, C, E, F.

## Discussion

Olig2 progenitor cells of the ventral spinal cord stop generating MNs and start producing OPCs in response to timely activation of Shh signaling. Here, we report that FGFs are key regulators of Shh-mediated fate change in Olig2 progenitor cells, acting both upstream of Shh and together with Shh to ensure specification of ventral OPCs (Figure 13). Our results support a model whereby newly differentiated ventral interneurons, that become providers of FGFs at the right time and place, might be initiators of the neuronal to oligodendroglial fate change in the ventral spinal cord.

**Figure 13:**
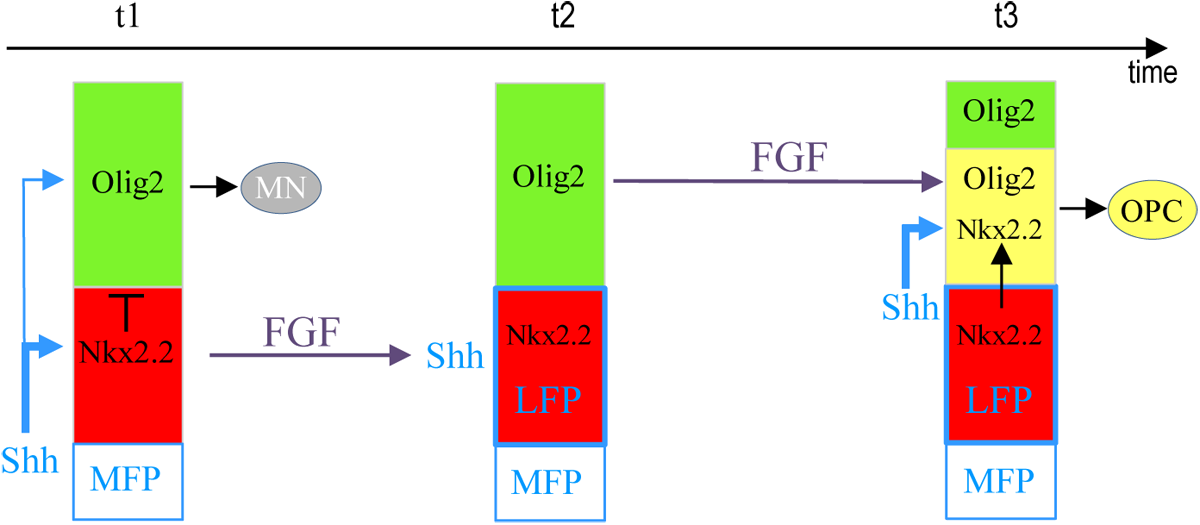
Model for FGF functions in OPC specification. At stage of ongoing neuronal generation (t1), the Olig2 (pMN, green) and Nkx2.2 (p3, red) progenitor domains do not overlap due to the repressive activity of Nkx2.2 on Olig2 expression. Olig2 progenitor cells generate motor neurons while Shh secreted by MFP cells activates low level of intracellular signaling (blue) to maintain expression of Olig2. As development proceeds (t2), Nkx2.2 progenitor cells become exposed to FGFs, possibly provided by ventral neurons. These cells subsequently up-regulate Shh expression to form the LFP. Soon after (t3), Olig2 progenitors, because they are submitted to higher Shh signalling activity, up-regulate Nkx2.2. At that time, Olig2 expression is no more repressed by Nkx2.2 thanks to joint exposure of these cells to FGFs. Olig2 and Nkx2.2 coexpressing progenitor cells, included in the p* domain (yellow), then start generating OPCs.

Shh-mediated signaling has long been recognized as essential for the ventral phase of oligodendrogenesis in the spinal cord. Because of its ability to promote OPC generation in the neural tube, Shh was also considered for some time to be sufficient to induce OPC commitment from neural progenitor cells (Trousse et al., 1995; Poncet et al., 1996; Pringle et al., 1996; Orentas et al., 1999; Soula et al., 2001; Danesin et al., 2006). The present study, by revealing that ventral OPC specification also needs FGFR signaling activation challenges this conclusion. Inactivation of FGFR signaling just prior to OPC specification proved to be as efficient to prevent generation of ventral OPCs as Shh signaling inhibition. The requirement of FGFR signaling for ventral spinal OPC specification is consistent with studies performed in various regions of the developing brain. Like in the spinal cord, the first wave of OPC production in the brain occurs in the ventral progenitor zone and requires Shh signaling activity (Alberta et al., 2001; Nery et al., 2001; Spassky et al., 2001; Tekki-Kessaris et al., 2001a; Fuccillo et al., 2004). However, evidence further emerged that lack of FGFR1/2 or FGFR inactivation in mouse and zebrafish result in failure of ventrally-derived brain OPCs to form properly (Esain et al., 2010; Furusho et al., 2011). Requirement of FGFR signaling for generation of ventral OPCs therefore comes up as a conserved process in the central nervous system. In agreement with Furusho and collaborators (2011), we found that FGFR signaling does not play a role in the regulation of cell proliferation or survival at stages of OPC generation but, instead, is part of the mechanism that triggers fate change of progenitor cells. We here highlight that FGFRs play a key role in changing the transcriptional identity of ventral progenitor cells over time, a change that ends up with induction of progenitor cells marked by Nkx2.2 and Olig2 coexpression. Activation of Nkx2.2 within Olig2 progenitor cells is well recognized to assign an OPC identity to these cells in chicken and zebrafish (Soula et al., 2001; Zhou et al., 2001; Agius et al., 2004; Kucenas et al., 2008; Al Oustah et al., 2014). Combined expression of Olig2 and Nkx2.2 is also efficient in triggering OPC specification from human embryonic stem cells (Hu et al., 2009). Although the role of Nkx2.2 in OPC specification is still uncertain in mouse, Nkx2.2 has been reported to be up-regulated in Olig2 progenitor cells at the time OPC specification also in this species (Sun et al., 2001; Touahri et al., 2012; Jiang et al., 2017) and this transcription factor, which is maintained in migratory OPCs, controls the timing of their differentiation (Qi et al., 2001; Sun et al., 2001; Fu et al., 2002; Gokhan et al., 2005; Cai et al., 2010; Zhu et al., 2014). Up-regulation of Nkx2.2 in Olig2 progenitor cells is known to depend on time activation of Shh signaling that occurs in the ventral spinal cord immediately prior to OPC specification (Danesin et al., 2006; Touahri et al., 2012; Al Oustah et al., 2014; Danesin and Soula, 2017). In agreement, Nkx2.2 is a direct but also high threshold Shh responsive gene (Lei et al., 2006; Vokes et al., 2007; Lek et al., 2010; Ribes et al., 2010). Our study, while not calling into question the role Shh in assigning an Nkx2.2 identity to Olig2 progenitor cells, reveals essential roles of FGFs in this process.

The present study unravels a complex interplay between Shh and FGFR signaling for ventral OPC induction. First, FGFR signaling, by inducing LFP formation, establishes a novel Shh signaling center in close proximity to Olig2 progenitor cells. FGFs therefore emerge as positive regulators of Shh expression in the ventral spinal cord. FGFs have already been involved in induction of Shh source cells, namely MFP cells (Sasai et al., 2014). However, mechanisms involved in induction of MFP and LFP are quite different. While FGFs regulate the competence of prospective MFP cells to respond to the Shh inductive signal (Sasai et al., 2014), we found that LFP induction in response to FGFR activation occurs independently of Shh. Importantly, failure of LFP induction, that occurs when FGFR are inactivated in LFP prospective cells of the p3 domain, is sufficient to prevent OPC specification. Therefore, Shh provided by MFP cells is not able to compensate the deficient production of Shh caused by lack of LFP formation. This is in agreement with our previous work showing that the extracellular enzyme Sulf1 contributes to induce OPCs by favoring provision of active forms of the morphogen by LFP cells but not by MFP cells (Touahri et al., 2012; Al Oustah et al., 2014). Thus, the present study not only directly demonstrates that LFP induction is an obligatory step to turn Olig2 progenitor cells to the OPC fate but also reveals the essential function of FGFs, acting upstream of Shh, in the induction of ventral OPCs.

In addition to their role in LFP induction, FGFRs proved to assume a second essential function in the control of OPC specification: they act cell-autonomously on progenitor cells of the pMN domain to ensure high level of Olig2 expression as these cells up-regulate Nkx2.2. Previous reports have convincingly shown that dorsal spinal progenitor cells in culture up-regulate Olig2 expression in response to FGFs (Chandran et al., 2003; Gabay et al., 2003; Kessaris et al., 2004; Cai et al., 2005; Abematsu et al., 2006; Bilican et al., 2008). However, different underlying molecular mechanisms, dependent of Shh (Gabay et al., 2003; Kessaris et al., 2004) or independent of Shh (Chandran et al., 2003; Cai et al., 2005; Abematsu et al., 2006; Bilican et al., 2008) have been proposed. The data presented here indicate that FGFR activation in spinal cord progenitor cells is not efficient in inducing Olig2 expression in progenitor cells in their endogenous context of the neural tube or when Shh signaling activity is impaired. Therefore, in the developing spinal cord, FGFR signaling regulates Olig2 expression at least in part through a Shh-dependent mechanism. Furthermore, at the time of OPC specification, attenuation of Olig2 expression by FGFR inactivation is noticed only when the LFP can form properly in ventral spinal cord explants. Olig2 expression in progenitor cells indeed appeared unaffected in experimental contexts where LFP formation has been prevented, i.e. when FGFRs were inactivated in both the p3 and pMN domains using FGFR inhibitors or when FGFR inactivation was limited to cells of the p3 domain by targeted overexpression of the dnFGFR. We therefore conclude that the cell-autonomous FGFR activation is needed to ensure maintenance of Olig2 expression only when these cells are submitted to high Shh signaling activity, i.e. when they activate Nkx2.2 expression. Reinforcing this conclusion, FGFR activation in the neural tube becomes a potent activator of Olig2 and Nkx2.2 coexpression when combined with Shh overproduction. Although the underlying mechanisms remains to be elucidated, this observation introduces FGF signaling as a way to prevent the Nkx2.2-dependent Olig2 repression in cells of the prospective p* domain as these cells upregulate Nkx2.2 in response to elevation of Shh signaling.

Two distinct models to account for FGF and Shh interplay in induction of ventral brain OPCs have been proposed. Esain and collaborators (2010) proposed a model in which Shh first establishes a progenitor cell domain competent to express Olig2 and then, FGFR signaling permits Olig2 gene transcription. By contrast, Furusho and collaborators (2011) proposed that FGFRs function downstream of Shh in the induction of ventrally derived OPCs. Our study by highlighting dual functions for FGFRs, acting both upstream of Shh and together with Shh to control OPC specification in the spinal cord might, somehow reconciles these apparently contradictory models. However, whether time activation of Shh signaling activity underlies initiation of OPC commitment in the developing brain remains to be established.

Our work opens the question of the source of FGFR ligands responsible for ventral OPC specification. Of the FGF members, FGF8, known to bind all four FGFRs and to control neural progenitor cell patterning in the embryonic brain (Crossley and Martin, 1995; Crossley et al., 1996; Ornitz et al., 1996), represents a good candidate. Populations of ventral spinal neurons, which progressively locate in close proximity to Olig2 progenitor cells, indeed up-regulate FGF8 at the right time to account for OPC induction. The interesting idea behind is that ventral spinal neurons, that are generated before OPC specification, might represent key initiators of oligodendrogenesis, an area that has yet to be explored.

In summary, we provide evidence that, in the ventral spinal cord, the coordination of Olig2 progenitor cell fate transition over time is primarily controlled by FGFR signaling. We furthermore elucidate an intimate interplay between FGFs and Shh in this process where FGFR activation is responsible for provision of high threshold Shh signal while ensuring the appropriate response to this signal to allow OPC specification.

## Acknowledgements

We acknowledge J. Guimera, S. Martinez, K. Storey, C. Tabin and R. Bansal who kindly provided us with valuable reagents. We are grateful to colleagues in the CBD for help, encouragement and advice, especially C. Danesin for helpful discussions and critical reading of the manuscript. We thank the Toulouse Regional Imaging Platform (TRI) for technical assistance in confocal microscopy. We acknowledge the Developmental Studies Hybridoma Bank, developed under the auspice of NICHD and maintained by the University of Iowa, Department of Biological Sciences, Iowa City, IA, for supplying monoclonal antibodies. Work in C.S.’ lab was supported by grants from ANR, ARSEP, CNRS and University of Toulouse. M-A.F. was supported by grant from French Ministry of National Education, Research, and Technology.

